# Identifying sub-populations of cells in single cell transcriptomic data – a Bayesian mixture modelling approach to zero-inflation of counts

**DOI:** 10.1101/2021.05.19.444841

**Authors:** Tom Wilson, Duong H.T. Vo, Thomas Thorne

## Abstract

In the study of single cell RNA-seq data, a key component of the analysis is to identify sub-populations of cells in the data. A variety of approaches to this have been considered, and although many machine learning based methods have been developed, these rarely give an estimate of uncertainty in the cluster assignment. To allow for this probabilistic models have been developed, but single cell RNA-seq data exhibit a phenomenon known as dropout, whereby a large proportion of the observed read counts are zero. This poses challenges in developing probabilistic models that appropriately model the data. We develop a novel Dirichlet process mixture model which employs both a mixture at the cell level to model multiple populations of cells, and a zero-inflated negative binomial mixture of counts at the transcript level. By taking a Bayesian approach we are able to model the expression of genes within clusters, and to quantify uncertainty in cluster assignments. It is shown that this approach out-performs previous approaches that applied multinomial distributions to model single cell RNA-seq counts and negative binomial models that do not take into account zero-inflation. Applied to a publicly available data set of single cell RNA-seq counts of multiple cell types from the mouse cortex and hippocampus, we demonstrate how our approach can be used to distinguish sub-populations of cells as clusters in the data, and to identify gene sets that are indicative of membership of a sub-population. The methodology is implemented as an open source Snakemake pipeline available from https://github.com/tt104/scmixture.

## 1 Introduction

Single cell transcriptomic data gives us a picture of the gene expression in individual cells as opposed to the measurements averaged over many cells produced by bulk experiments. A particular methodology known as single cell RNA-seq (scRNA-Seq) has grown massively in capacity and popularity in recent years (Svensson et al. 2018). This approach allows for the heterogeneity in populations of cells to be explored by identifying sub populations of cells based on the measured expression of transcripts in individual cells. This is especially relevant in tissues where it is not physically possible to extract cells of a specific known type, for example in the brain (Lake et al. 2018, Mathys et al. 2019), where distinct cell types can be inferred from data generated from a mixture of cells of different types. The challenge in this scenario is to allow cell types within a sample to be identified as an unsupervised learning problem, where we wish to cluster together cells of similar types, based on their expression (Stegle et al. 2015). Although this can be approached by mapping cells to specific reference expression patterns, there will always be scenarios where typical expression patterns are not available (Szabo et al. 2019), or where cells with atypical expression are of interest, for example in disease studies (Mathys et al. 2019, Liu et al. 2021).

Work in the literature has applied a range of different approaches to this problem, some making use of probabilistic methods, whilst others apply algorithmic machine learning techniques, and some use a mixture of the two. Probabilistic model based approaches include the use of a multinomial model of counts derived from single cell experiments in a Dirichlet process mixture model (Duan et al. 2019), and hierarchical Bayesian nonparametric models which allow for clustering of data at both the cell and subject level (Wu & Luo 2021). These Bayesian nonparametric approaches have the advantage that the number of components in the model is learnt from the data, and we adopt a similar Dirichlet process mixture modelling approach in this work. In Duan et al. (2019) the single cell RNA-seq data are modelled as counts following a multinomial distribution, as this circumvents issues around zero-inflation, but the multinomial distribution may not provide a good fit to scRNA-seq count data, and the (zero-inflated) negative binomial distribution produces a significantly better fit, as discussed below. A benefit of the approach in Duan et al. (2019) is that the implementation can be run in parallel over multiple cores on an HPC cluster, and it is demonstrated that this can greatly improve performance when using larger data sets. In Wu & Luo (2021) a more realistic zero-inflated negative binomial model of scRNA-seq count data is applied, but the model is designed to tackle the more complex problem of clustering at the level of both subjects and cells for experimental data generated from multiple subjects. This applies a hybrid hierarchical Dirichlet process and nested Dirichlet process model to represent the hierarchical structure in the data, and can learn the number of subject groups and cell groups from the data.

Some approaches, for example Monocle (Trapnell et al. 2014), SC3 (Kiselev et al. 2017), and Seurat (Stuart et al. 2019) apply clustering techniques such as K-means, DBSCAN or Louvain clustering, after applying transformations to the data to produce a representation of each cell more amenable to these approaches (Zhang et al. 2020). While the clustering itself is then often fast and straightforward to apply, such methods do not attempt to model the statistical distribution of the data, and unlike Bayesian nonparametric methods, often require a trial and error approach to parameter tuning, for example applying the model to each possible number of clusters in the data and applying heuristics to determine the number of clusters in the data. However these methods are often still faster to apply than more complex models described above that rely on Markov Chain Monte Carlo (MCMC) sampling.

Several machine learning based approaches applying neural networks using autoencoders or similar embedding approaches have been proposed (Lopez et al. 2018, Grønbech et al. 2020, Svensson et al. 2020), representing expression data in a lower dimensional vector space, on which clustering using methods such as K-means can be performed. However neural network based approaches such as autoencoders suffer from the drawback that results are not stable between different runs unless the random seed is fixed, and so multiple autoencoder neural networks trained on the same data will give different embeddings, with no clear indication of which are superior. This criticism could also be made of Bayesian probabilistic models, but it is possible to select a best clustering from the results of multiple runs based on the posterior probability. There is also no quantification of uncertainty in cluster assignments using neural network based approaches. This lack of quantification of uncertainty and its further propagation through subsequent stages of an analysis pipeline has been identified as a key challenge it the future development of single cell data analysis pipelines (Lähnemann et al. 2020).

A particular benefit of a probabilistic approach is in the interpretability of the model, in terms of the ability to link certain parameters of the model directly to properties of the features in the data. As a result the learned clusters can easily be associated with patterns of gene expression through the model. Although attempts have been made to build machine learning models that are interpretable in this sense, for example applying autoencoders with linear decoders (Svensson et al. 2020), these methods still lack the inclusion of posterior uncertainty provided by Bayesian models, both in terms of cluster assignment and gene expression within clusters, and still suffer from the issues with reproducibility described above.

Recently latent factor models have also been applied to scRNA-seq data (Levitin et al. 2019), and allow us to identify important patterns in gene expression across cells. Although these methods can produce useful biological insights, and they are capable of identifying factors that correspond to individual cell types, the task they perform is not directly related to that of clustering. Rather than identifying patterns of expression that define distinct groups of cells, latent factor models learn a set of basis expression signatures that can be combined in different weights to represent the expression across all cells in the data, and for the model used in Levitin et al. (2019), the number of factors to learn must be specified in advance. As a result this method is less useful for the specific task of identifying sub-populations of cells within a data set.

Single cell transcriptomic data has been said to suffer from a phenomenon know as dropout. The occurrence of dropout events can be due to three main reasons (Qiu 2020), the stochastic expression of genes which has been demonstrated by multiple studies and experiments (Raj & van Oudenaarden 2008), the low expression of genes in individual cells which is demonstrated through the detection of low-expression gene TERT by only qPCR in the study of Wu et al. (2014), and the inefficiency of mRNA capture and amplification (Haque et al. 2017). The result is the large number of zero counts in scRNA-seq data, which in turn leads to distributions of counts with large spikes at zero. There has been some discussion in the literature around the issue of zero inflation in single cell transcriptomic data, and it is argued in Svensson (2020) that the proportions of zero counts are no larger than would be expected from a negative binomial distribution in data collected from droplet based sequencing platforms. However in Clivio et al. (Clivio et al. 2019) a Bayesian model selection approach demonstrates that this is not the case across all data sets, even in protocols which use unique molecular identifiers (UMIs), and that the phenomenon may only be observed in a subset of genes. Previous work in the literature has used probabilistic models of zero-inflation to model dropout in scRNA-seq data, including scVI (Lopez et al. 2018), ZINB-WaVE (Risso et al. 2018), and scVAE (Grønbech et al. 2020). The ZINB-WaVE approach applies a flexible zero-inflated negative binomial model to dimensionality reduction of scRNA-seq data (Risso et al. 2018), that allows for dependence on gene expression and other covariates in the dropout rate. It was found that ZINB-WaVE produced more biologically relevant low dimensional representations of scRNA-seq data than PCA as it was able to remove unwanted sources of variation in the data. scVI and scVAE both apply variational autoencoders to generate a latent representation of the cells in scRNA-seq data, and use zero-inflated negative binomial distributions to model the data. In Lopez et al. (2018) it was found that models that applied a zero-inflated negative binomial distribution to model the data performed better than those that do not.

Interestingly, previous work in the literature (Qiu 2020) has also shown that taking into account the pattern of dropout alone provides enough information to separate cell types in scRNA-seq data. It is hypothesised in Qiu (2020) that genes in the same pathway tend to exhibit the same pattern of dropout over cell types, so that a cell type can potentially be identified by the pattern of dropout across cells.

To investigate the use of zero-inflated negative binomial models for scRNA-seq data, we considered multiple data sets generated from different protocols and applied a straightforward model selection approach to counts within cell types. We found that a zero-inflated negative binomial distribution is a superior model for a small percentage of genes across multiple experimental platforms, and that the dropout rate for a single gene varies between cell types. These results are presented in the supplementary materials.

To represent dropout in scRNA-seq data in a probabilistic clustering model, we have developed a Bayesian nonparametric clustering method that employs a zero-inflated negative binomial model with a transcript and cell type dependent mixing proportion. This allows genes that do exhibit inflated zero counts to be appropriately modelled, and is outlined in the methods section below.

## 2 Methods

We model the expression of each transcript as following a zero-inflated negative binomial distribution, with parameters defining the mean, dispersion, and proportion of dropout. For each cell *i* and each transcript *j*, counts of reads *y*_*ij*_ are modeled as a mixture of an atom at zero, representing dropout, and a negative binomial distribution with mean *μ*_*j*_, dispersion *ω*_*j*_ and transcript specific dropout rates *w*_*j*_,

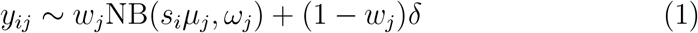

where *δ* is the Dirac delta distribution. There is also a fixed cell specific scaling factor *s*_*i*_ that compensates for the variation in sequencing depth between cells. Scaling factors can be estimated using existing approaches in the literature, for example scran (Lun et al. 2016).

### 2.1 Bayesian nonparametric clustering model

We introduce a clustering of the data to represent sub-populations of cells by placing a Dirichlet Process prior DP(*α, H*) with concentration parameter *α* on the mean, dispersion and dropout rate parameters of all genes. Here the base measure *H* corresponds to a prior on *μ, ω* and *w*. The Dirichlet process prior can be represented using the stick breaking construction of the Dirichlet process (Sethuraman 1994), which introduces stick breaking proportions *u*_*l*_, and stick lengths *β*_*k*_ which depend only on the *u*_*l*_, for a potentially infinite number of *u*_*l*_ and *β*_*k*_. Sampling is performed by introducing a latent variable describing the cluster membership of each cell, *z*_*i*_. Then we can write:

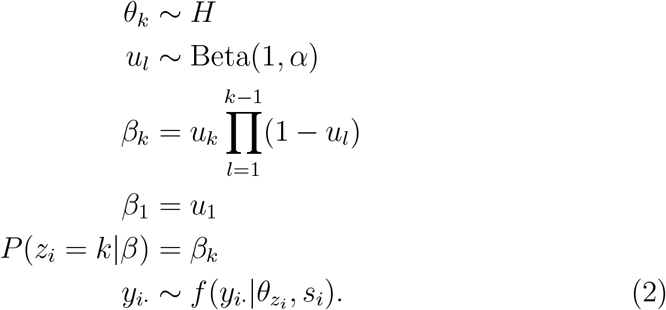

where *H* is the prior on *μ, ω* and *w, θ*_*k*_ = (*μ*_*k*·_, *ω*_*k*·_, *w*_*k*·_), the cluster specific mean and dispersion and zero-inflation parameters for each gene, and *k* and *l* range over ℤ ^+^. The model is illustrated as a graphical model in figure 1.

**Figure 1:**
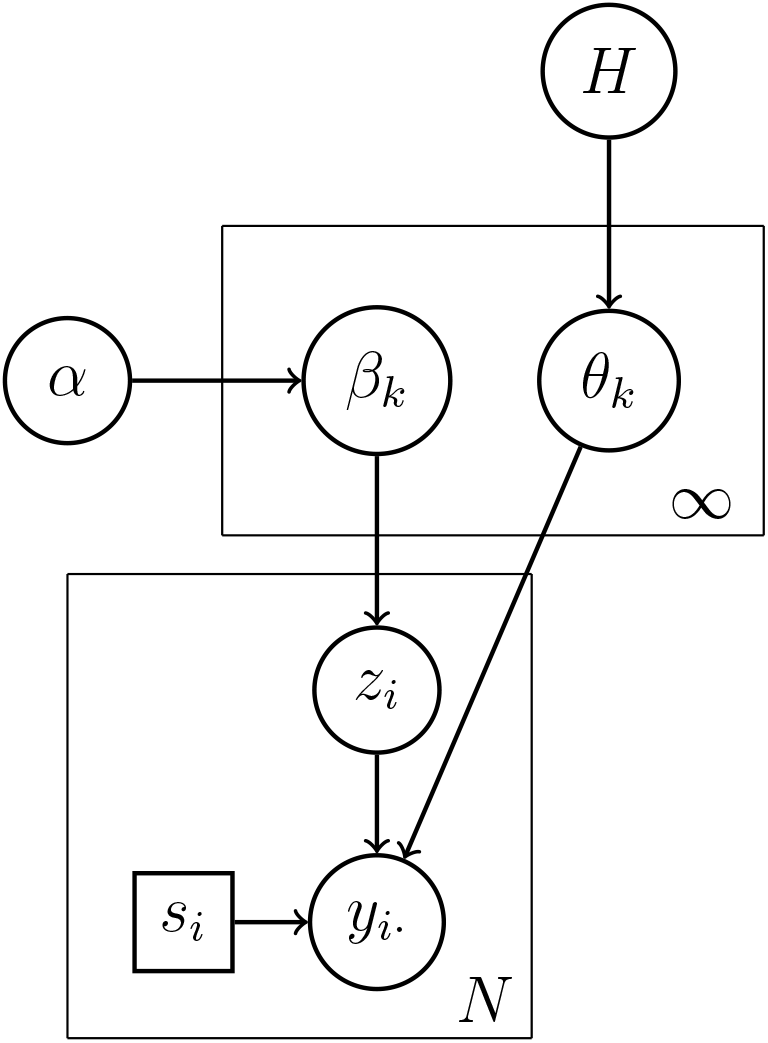
A graphical model representation of the Dirichlet process mixture. The parameter *θ* represents the mean *μ*_*k*·_, dispersion *ω*_*k*·_ and dropout rate *w*_*k*·_ for cluster *k*.

For the base measure *H* of the Dirichlet process which specifies the priors on the cluster specific values of *θ*_*k*_ = (*μ*_*k*·_, *ω*_*k*·_, *w*_*k*·_) we use Gamma distributions for *μ* and *ω* and a uniform Beta(1, 1) prior for *w*. To set the parameters of the Gamma distributions we use an empirical Bayes approach, whereby we estimate the distributions of *μ* and *ω* across all genes by fitting the model in equation 1 to each gene. We do this using maximum likelihood inference to estimate values 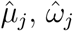 and 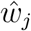 for each gene *j*. Maximum likelihood inference was performed using the L-BFGS-B bounded numerical optimisation algorithm as implemented in the optim function in the R statistical environment.

This produces a set of estimates of 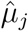 and 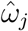 for *j* ∈1, …, *P*, where *P* is the total number of transcripts. We can then estimate the distribution of *μ* and *ω* across all transcripts by using maximum likelihood inference to fit Gamma distributions to the sets of values 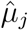 and 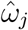, again using numerical optimisation. We parameterise the component of *H* representing the prior on *μ*_*k*_ as Gamma(*a, b*) and the prior on *ω*_*k*_ as Gamma(*s, t*), with the values of *a, b, s* and *t* determined using the process described above.

### 2.2 Markov Chain Monte Carlo sampling scheme

To sample the cluster allocation variables *z*_*i*_ and cluster specific parameters *μ*_*k*·_, *ω*_*k*·_ and *w*_*k*·_, we use the stick breaking sampler described in Hastie et al. (2014) and Liverani et al. (2015). This samples variables of the stick breaking construction *u*_*l*_ through the use of auxiliary variables, and uses retrospective sampling to sample parameters for unoccupied clusters, and then samples each *z*_*i*_ conditionally on the stick breaking lengths. The sampling scheme is outlined in algorithms 1, 2 and 3.

#### Algorithm 1

MCMC sampler algorithm

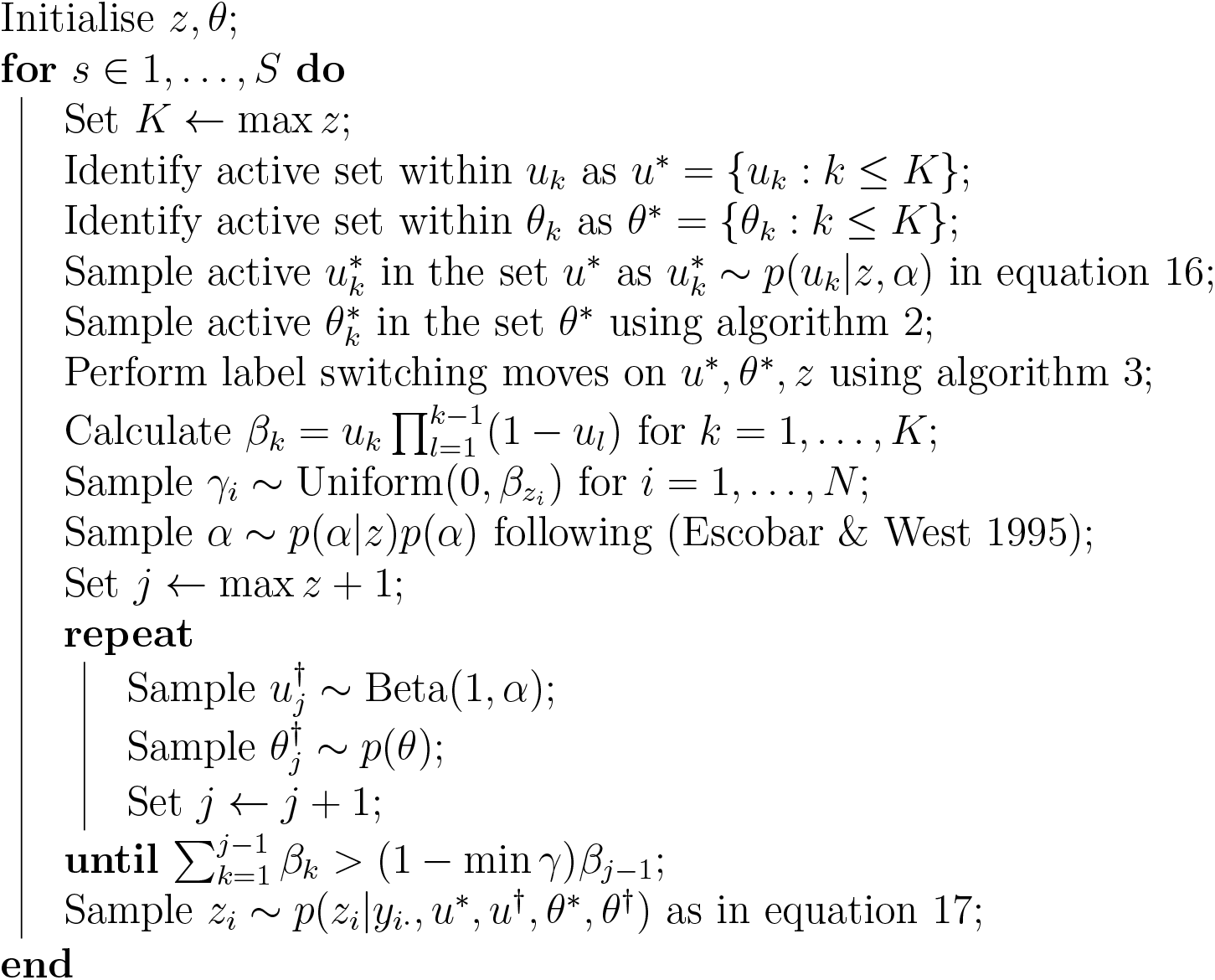

To derive MCMC updates of the cluster specific parameters *μ*_*kj*_ and *ω*_*kj*_ we can use Gibbs sampling steps whereby each parameter value is updated by drawing a sample from the conditional distribution of the parameter given the remaining parameters. To allow us to sample from the posterior on the cluster specific mean and dispersion parameters of the negative binomial distribution for cluster *k, μ*_*kj*_ and *ω*_*kj*_, we introduce auxiliary variables *v*_*ij*_ to model the negative binomial distribution as a Poisson–Gamma mixture. This gives

#### Algorithm 2

MCMC sampler for active clusters

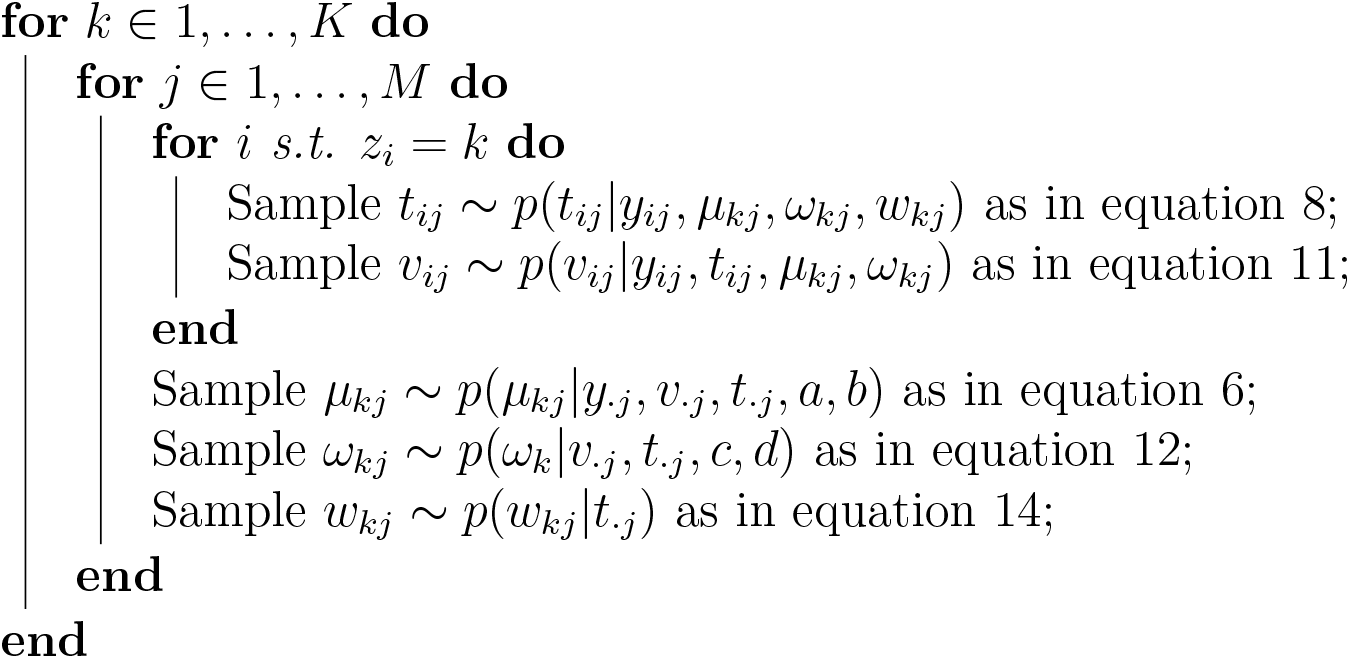

#### Algorithm 3

Label switching moves

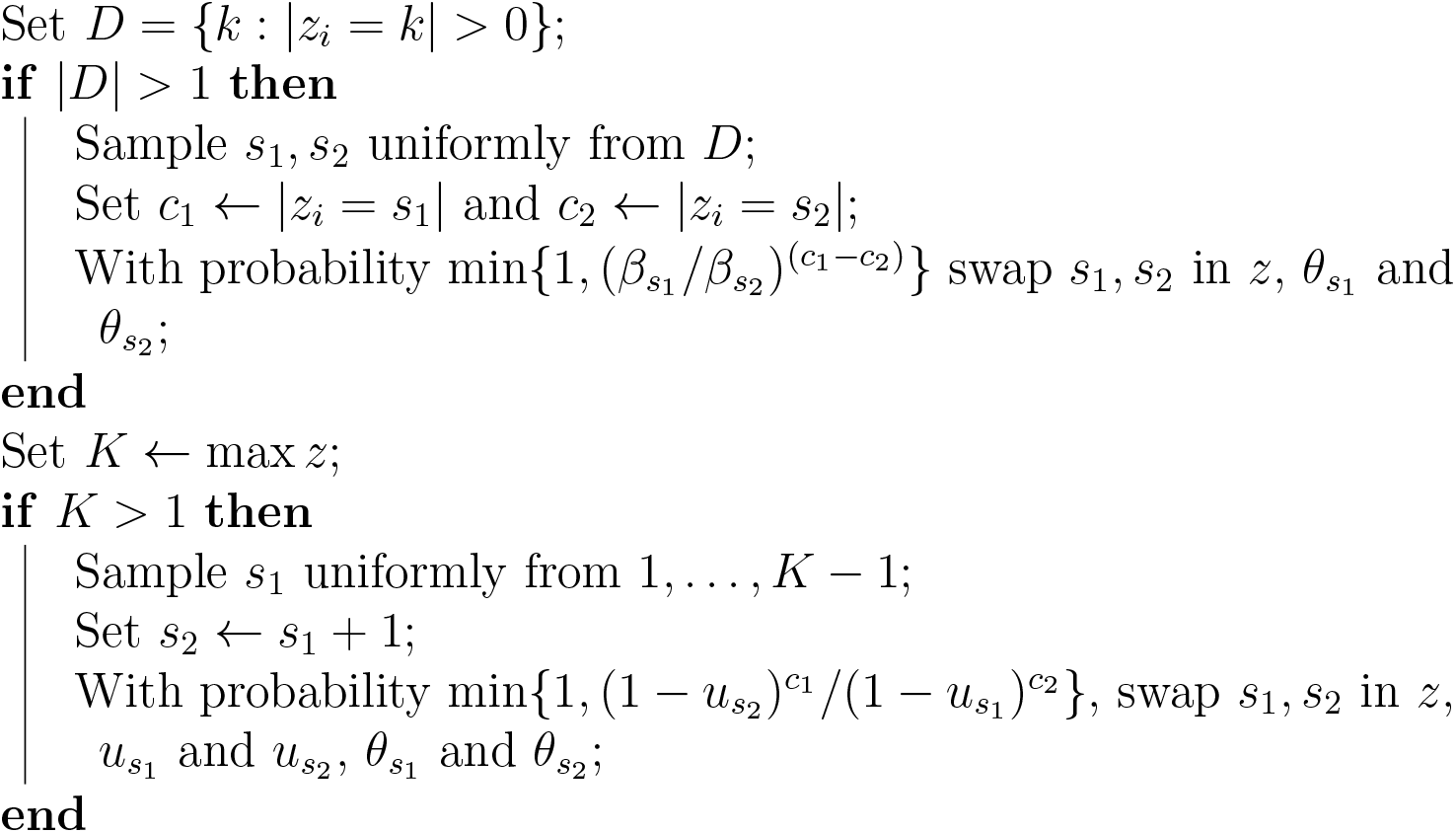

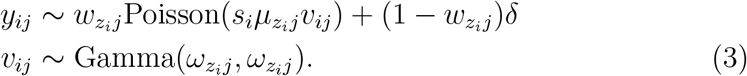

The cluster specific mixture weights *w*_*kj*_ in equation 3 are sampled by introducing latent indicator variables *t*_*ij*_ so that

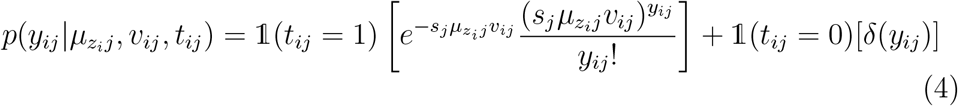

For each transcript *j* within a cluster *k*, there is a cluster specific mean *μ*_*kj*_, with prior Gamma(*a, b*) that is sampled conditionally in a Gibbs sampling update step as

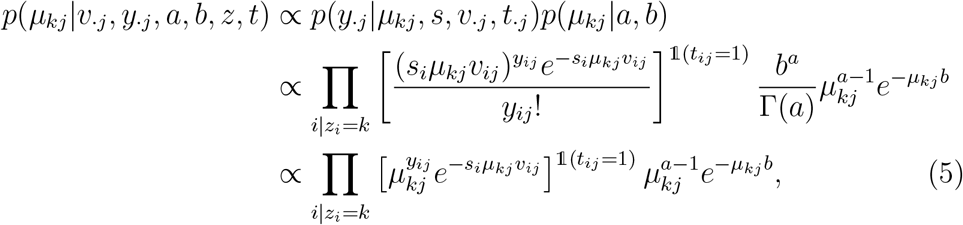

collecting terms that vary with *μ*_*kj*_ in equation 5 and reinstating the normalising constant gives a Gamma distribution:

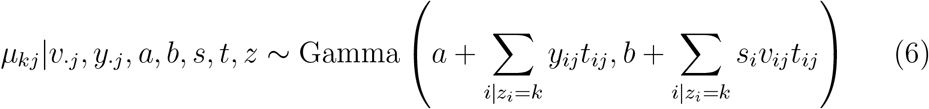

The Gibbs sampling update for the indicator *t*_*ij*_ is derived using

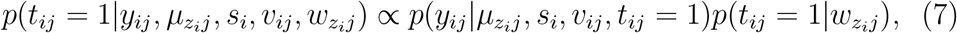

and similarly for *t*_*ij*_ = 0, so *t*_*ij*_ can be updated by sampling from a binomial distribution with

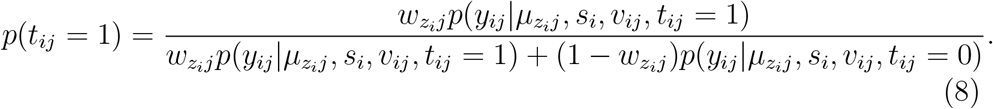

using equation 4 for 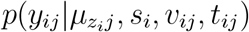.

Then given *t*_*ij*_ = 1 we can use Gibbs sampling for *v*_*ij*_ as needed, for *i* such that *z*_*i*_ = *k* using

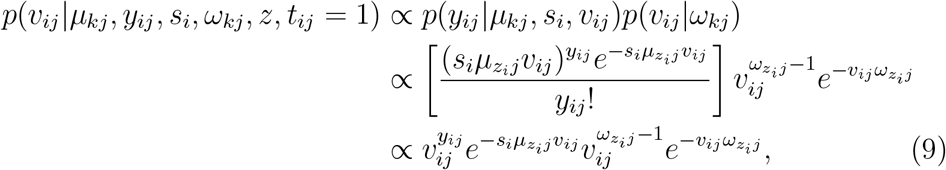

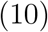

where collecting terms in *v*_*ij*_ in equation 10 we can observe that

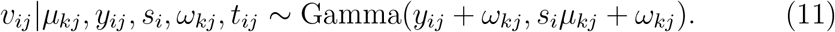

To sample the dispersion parameter *ω*_*kj*_, using the definition in equation 3 and with prior Gamma(*c, d*) we apply a slice sampler (Neal 2003) on *p*(*ω*_*kj*_|*v*_·*j*_, *c, d, t, z*) for each *j* independently:

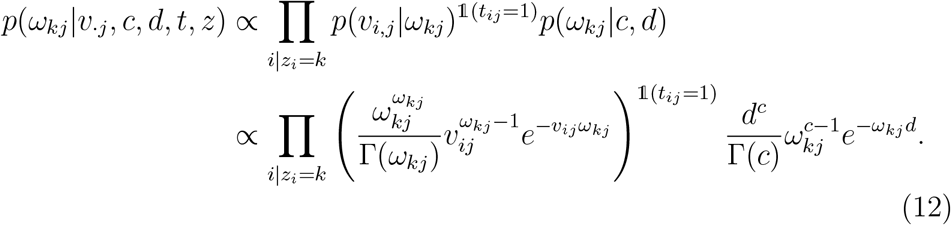

Then cluster specific weights *w*_*kj*_ for zero-inflation are sampled using the *t*_*ij*_ as

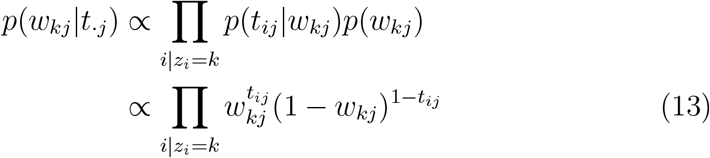

and so we can update *w*_*kj*_ by sampling from

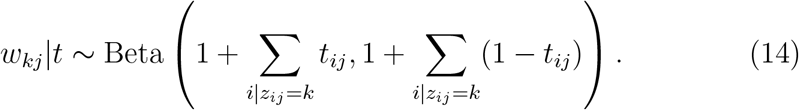

The MCMC sampler is outlined in algorithm 1, and iterates over a loop that updates the within cluster specific parameters *μ*_*k*·_, *ω*_*k*·_, and *w*_*k*·_ sampling the appropriate *v*_*ij*_ and *t*_*ij*_ as needed to do so as described in algorithm 2, and updates the cluster allocation variables *z*_*i*_ using the sampling scheme described in Hastie et al. (2014) and Liverani et al. (2015). This uses auxiliary variables *γ*_*i*_, as originally introduced in Walker (2007), so that

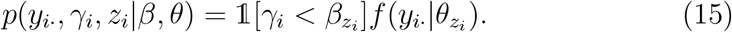

The sampler defines sets of active (*u*^*^ and *θ*^*^) and potential (*u*^†^ and *θ*^†^) variables *u* and *θ*, based on the *γ*_*i*_. Conditional on *z* and *α* the active *u*^*^ are updated using

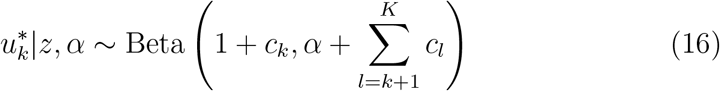

where *c*_*k*_ = |*z*_*i*_ = *k*| and *K* = max *z*. The label switching moves of Papaspiliopoulos & Roberts (2008) are used on *u*^*^, *θ*^*^ and *z* to improve mixing, as described in algorithm 3. To sample *u*^†^ in algorithm 1, the sampler loops over increasing *j >* max *z* sampling 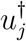 until 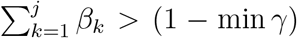. The Dirichlet process concentration parameter *α* is given a Gamma(1, 1) prior and updated using the auxiliary variable Gibbs sampling scheme of Escobar & West (1995). Given *u* and *θ*, it is then possible to sample each *z*_*i*_ from a multinomial using

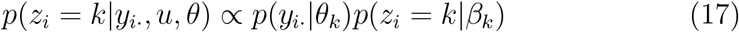

where the index *k* ranges over all indices in *u*^*^, *u*^†^ and *θ*^*^, *θ*^†^, and *p*(*y*_*i*·_|*θ*_*k*_) is as in equation 4.

## 3 Results

We applied our clustering method to data from benchmarking experiments (Tian et al. 2019) as well as an scRNA-Seq data set from the literature (Zeisel et al. 2015). For comparison, we implemented a model where the counts within a cluster follow a multinomial distribution, in the same Dirichlet process mixture model sampling framework as used in our zero-inflated negative binomial model. Scaling factors for each cell were inferred using the scran (Lun et al. 2016) R package. As in other probabilistic scRNA-Seq data analysis methods (Duan et al. 2019, Wu & Luo 2021), to reduce the computational burden we first filtered the set of transcripts, taking the 1000 with the highest coefficient of variation across all cells when applying the multinomial and negative binomial models. The cluster allocations in the MCMC samplers were initialised uniformly at random to 100 different clusters. We ran four Markov Chain Monte Carlo (MCMC) chains with burn-in for 1000 iterations of the outer loop of the sampler, and selected the maximum a posteriori (MAP) estimates of the allocation variables *z*_*i*_ across all samples collected to avoid issues with label switching (Jasra et al. 2005). Convergence was assessed using the Gelman-Rubin diagnostic (Gelman & Rubin 1992) on *α*. We compare our method to an implementation using a multinomial count distribution, similar to that used in Duan et al. (2019), an implementation using a negative binomial count distribution without zero-inflation, and two of the top performing scRNA-Seq clustering algorithms studied in Tian et al. (2019), SC3 (Kiselev et al. 2017) and Seurat (Stuart et al. 2019). Both SC3 and Seurat were run on the log transformed data using default settings.

### 3.1 Single cell benchmarking data

To test the performance of our method on data where there is a known ground truth set of cell populations, we applied our methodology to the single cell benchmarking data sets of Tian et al. (2019). These provide data from various mixtures of human lung adenocarcinoma cell lines, designed for the benchmarking of single cell RNA-Seq analysis pipelines, and were sequenced using a number of different platforms.

The first two benchmarking data sets, the three cell and five cell mixtures, are comprised of mixtures of cells from HCC827, H1975, and H2228 cell lines, and mixtures of cells from HCC827, H1975, H2228, A549, and H838 cell lines respectively. These were sequenced using the plate based CEL-seq2 (Hashimshony et al. 2016) platform, and the droplet based 10x (Zheng et al. 2017) and Drop-seq (Macosko et al. 2015) platforms.

The different methods considered were evaluated using the Adjusted Rand Index (ARI)(Hubert & Arabie 1985) using the known cell type labels included in the data, over multiple repeated runs of the methods on the data. The Rand index measures the similarity between two sets of cluster labels for the data, and the ARI adjusts this to use the expected performance of a random clustering assignment as a baseline. Using default settings for SC3 and Seurat the results did not vary between runs as the random seed is fixed in the default parameter values. However to illustrate the potential variability in performance we ran the methods multiple times with different random seeds, as this gives a more accurate representation of how performance may vary across data sets. The multinomial and zero-inflated negative binomial models use MCMC methods to sample from a model posterior, and so are inherently subject to some variation in the results between runs.

Results from the three cell mixture are shown in figure 2(a). It can be seen that our negative binomial model with zero-inflation outperforms the multinomial and negative binomial models and the SC3 and Seurat methods in the 10x and Dropseq data sets, and is second in performance in the CEL-seq2 data set. As illustrated in figure 2(c), performance is similar in the five cell mixture data sets, with the negative binomial model with zero-inflation providing a clear advantage over the multinomial and negative binomial models in all cases, whilst performing similarly to Seurat and SC3. Interestingly, even using a varying random seed, the performance of Seurat is highly stable across runs.

**Figure 2:**
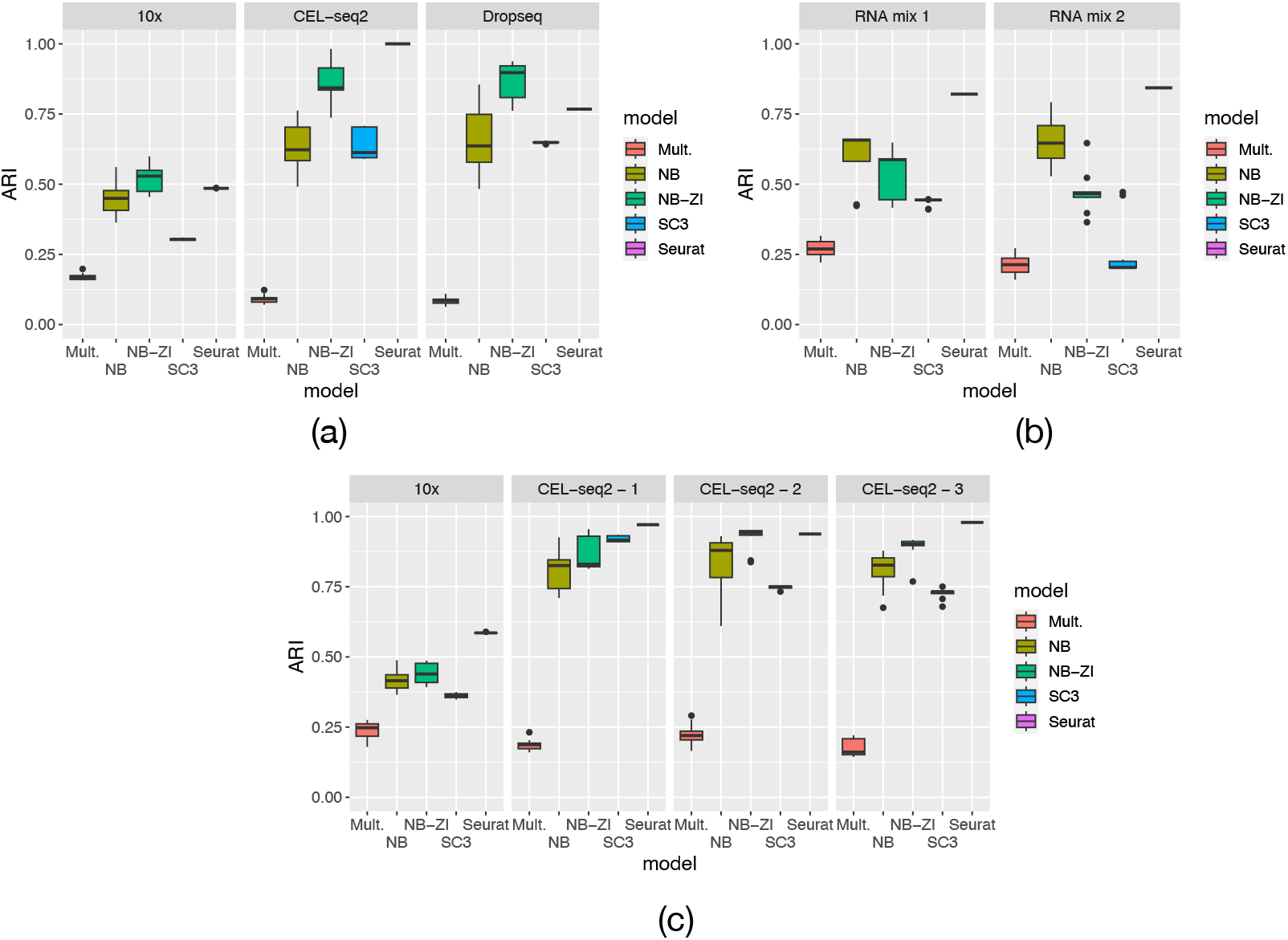
Performance measured by Adjusted Rand Index for 10 runs of each clustering model, across multiple data sets – (a) a mixture of 3 cell types across multiple experimental platforms, (b) a mixture of RNA in 7 different proportions from bulk cell populations for 3 different cell types (c) a mixture of 5 cell types across several experimental platforms.

Another benchmarking sample in Tian et al. (2019) is constructed as a mixture of bulk RNA from three different cell lines, combined in different proportions to produce seven different pseudo cells. Results comparing the performance of the methods we consider are shown in figure 2(b). Compared to the three and five cell mixtures, the performance of the negative binomial model is improved over the zero-inflated version. We hypothesize that the improvement in the negative binomial model is due to the artificial construction of the samples used to produce the data. Specifically by producing pseudo cells through effectively summing bulk RNA quantities from different cell lines, the data are no longer counts from single cells, but rather a weighted sum of counts from multiple cell types.

We also consider the number of clusters learnt by each model, with the results illustrated in figure 3. It can be seen that across all three of the bench-marking data sets, our zero-inflated negative binomial approach is closer to identifying the correct number of clusters than the multinomial model, which significantly overestimates the number of clusters. For the three cell and five cell mixtures sequenced on the 10x platform, it can be seen that in terms of identifying the number of clusters, our method outperforms SC3, while performing similarly to SC3 and Seurat on CEL-seq2 and Dropseq data. Overall it can be seen that Seurat is the most reliable method in estimating the correct number of clusters from the data, and outperforms both our approach and SC3.

**Figure 3:**
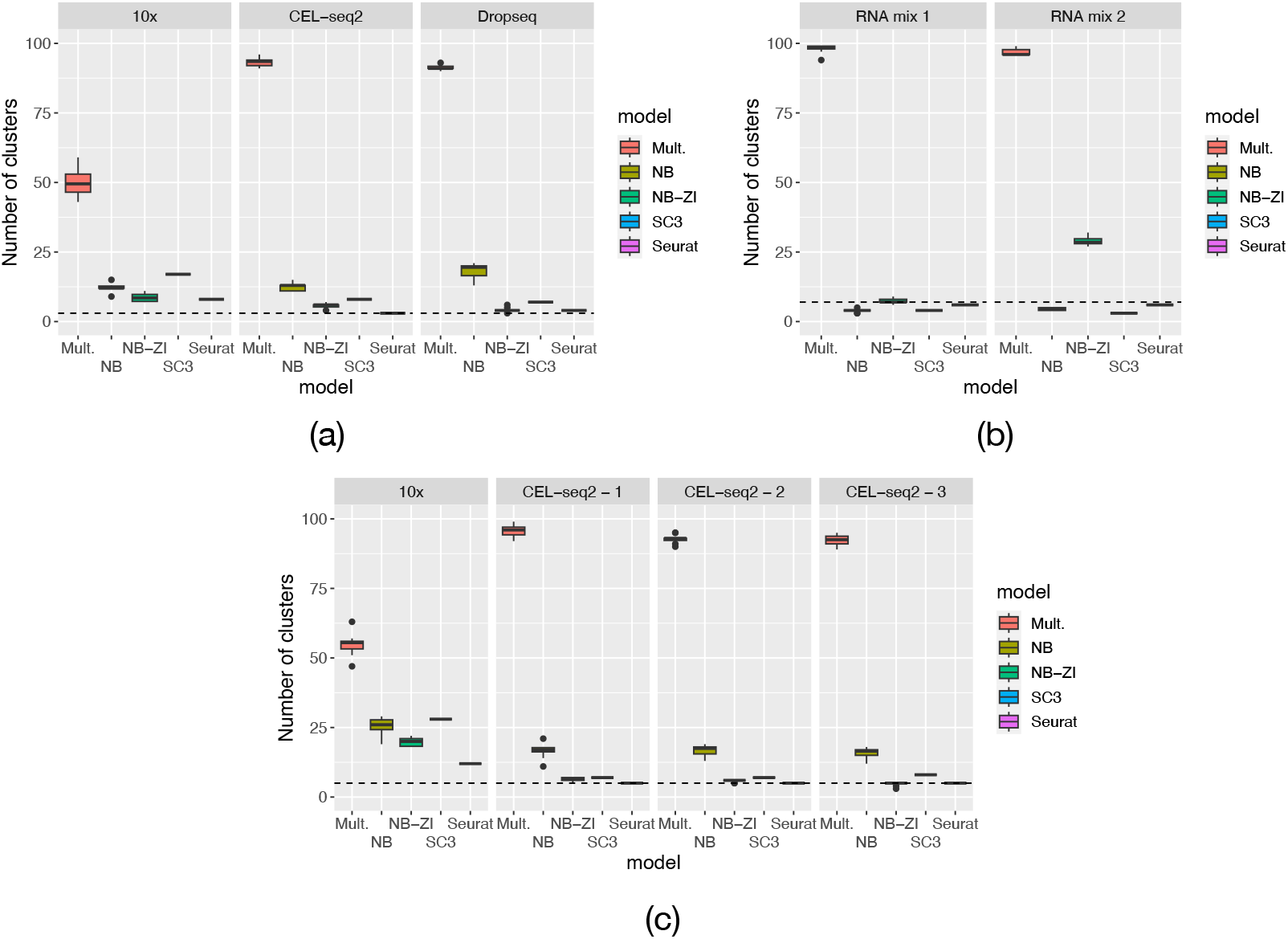
Numbers of clusters inferred (horizontal lines indicate true cluster number) for 10 runs of each clustering model across multiple data sets – (a) a mixture of 3 cell types across multiple experimental platforms, (b) a mixture of RNA in 7 different proportions from bulk cell populations for 3 different cell types (c) a mixture of 5 cell types across several experimental platforms.

### 3.2 Mouse cortex and hippocampus single cell data

To illustrate the results that we can produce using our model, our methodology was applied to the single cell RNA-Seq data set of Zeisel et al. (2015), that provides data for 3005 cells from the mouse cortex and hippocampus. In Zeisel et al. (2015) a clustering produced using the BacksSPIN algorithm, a bi-clustering based variant of the SPIN algorithm (Tsafrir et al. 2005), is refined using heuristic approaches to produce the clustering shown in figure 4. Applying our method to the data produces 23 clusters, many of which are clearly aligned to the cell types identified in the data in Zeisel et al. (2015). In some cases our clustering subdivides the higher level cell type specific clusters identified by Zeisel et al., who also observed lower level subdivisions of cell types into further clusters. We also include the clusterings produced by Seurat and SC3 on this data in figure 4. It can be seen that both methods also produce clusters with a clear correspondence to the annotation from Zeisel et al. (2015), although SC3 produces a much larger number of clusters.

**Figure 4:**
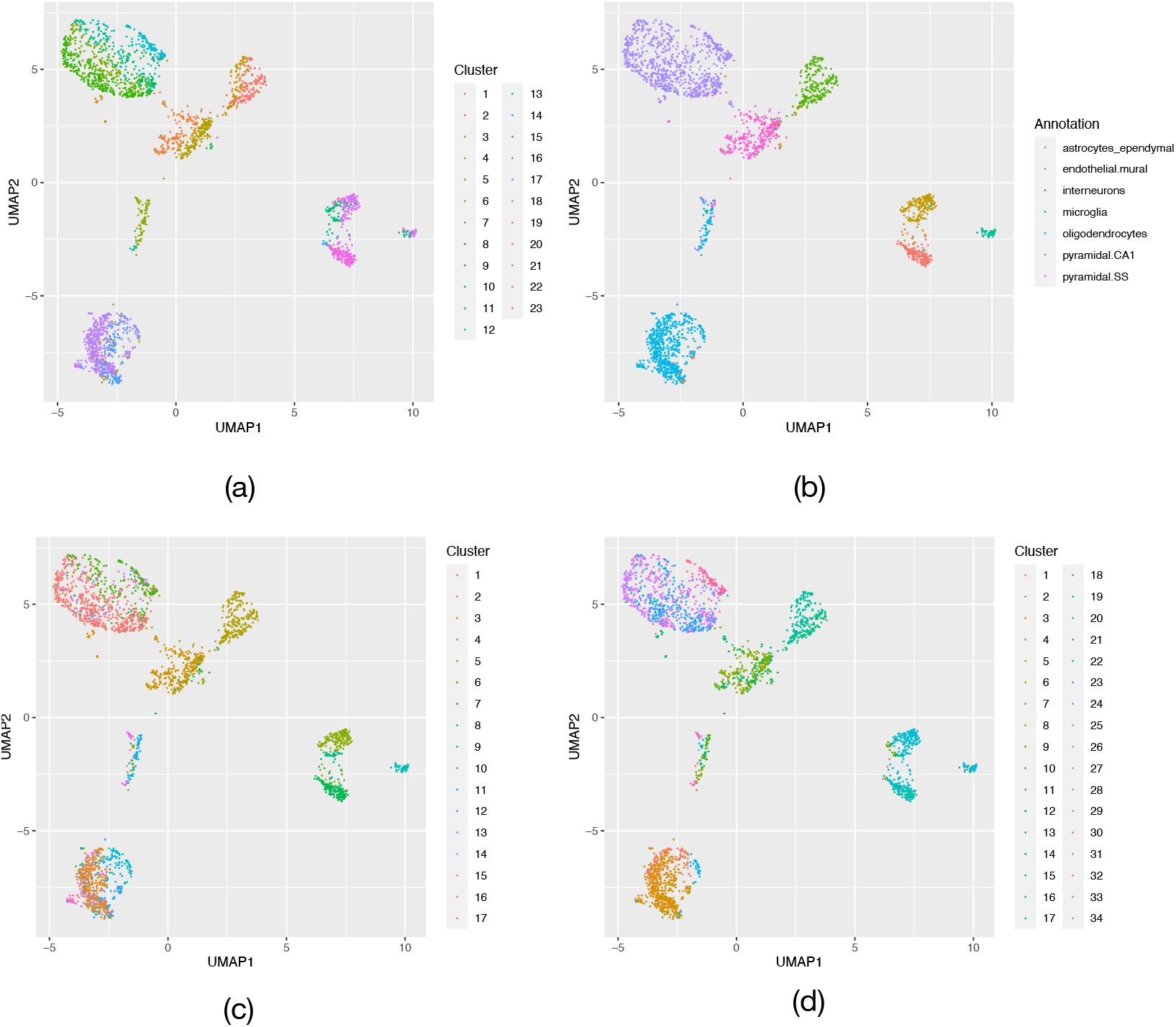
UMAP embeddings of cells from the mouse cortex and hippocampus data with (a) clusters generated by our zero-inflated negative binomial model, (b) annotations from Zeisel et al. (2015), (c) a clustering generated by Seurat and (d) a clustering generated by SC3.

A benefit of our approach is that we can investigate the MAP estimates of the mean and dispersion parameters for each cluster. This allows us to easily identify key genes that are important in defining cluster membership. We do so by comparing the mean expression of the gene in a cluster with the average expression across all clusters, taking into account the mean and dispersion of the counts. Genes whose mean expression level in a cluster is above the 95th percentile or below the 5th percentile of the distribution of the across-cluster average count distribution are then taken as being up or down regulated in the cluster.

We are also able to remove cells whose cluster membership cannot be identified with confidence. To do so we calculate the mean posterior probability of a cell belonging to the same cluster as the other cells in its MAP estimated cluster. Then we can filter out cells where this mean probability is below a certain threshold, here set to 0.5.

A UMAP (McInnes et al. 2020) representation of the data combined with our clustering output is shown in figure 4. We focus on four specific clusters, 1, 7, 18, and 20 highlighted in figure 5, which are identified as interneurons, pyramidal CA1 cells, oligodendrocytes and astrocytes respectively in Zeisel et al. (2015). Considering the inferred gene expression in these clusters, our method identifies a set of overexpressed genes in all four. The cluster specific means of genes in clusters 1, 7, 18 and 20 are illustrated in figure 6. Using g:Profiler (Raudvere et al. 2019) we also identified enriched biological pathways of upregulated genes from the KEGG (Kanehisa & Goto 2000) and Reactome (Gillespie et al. 2022) databases.

**Figure 5:**
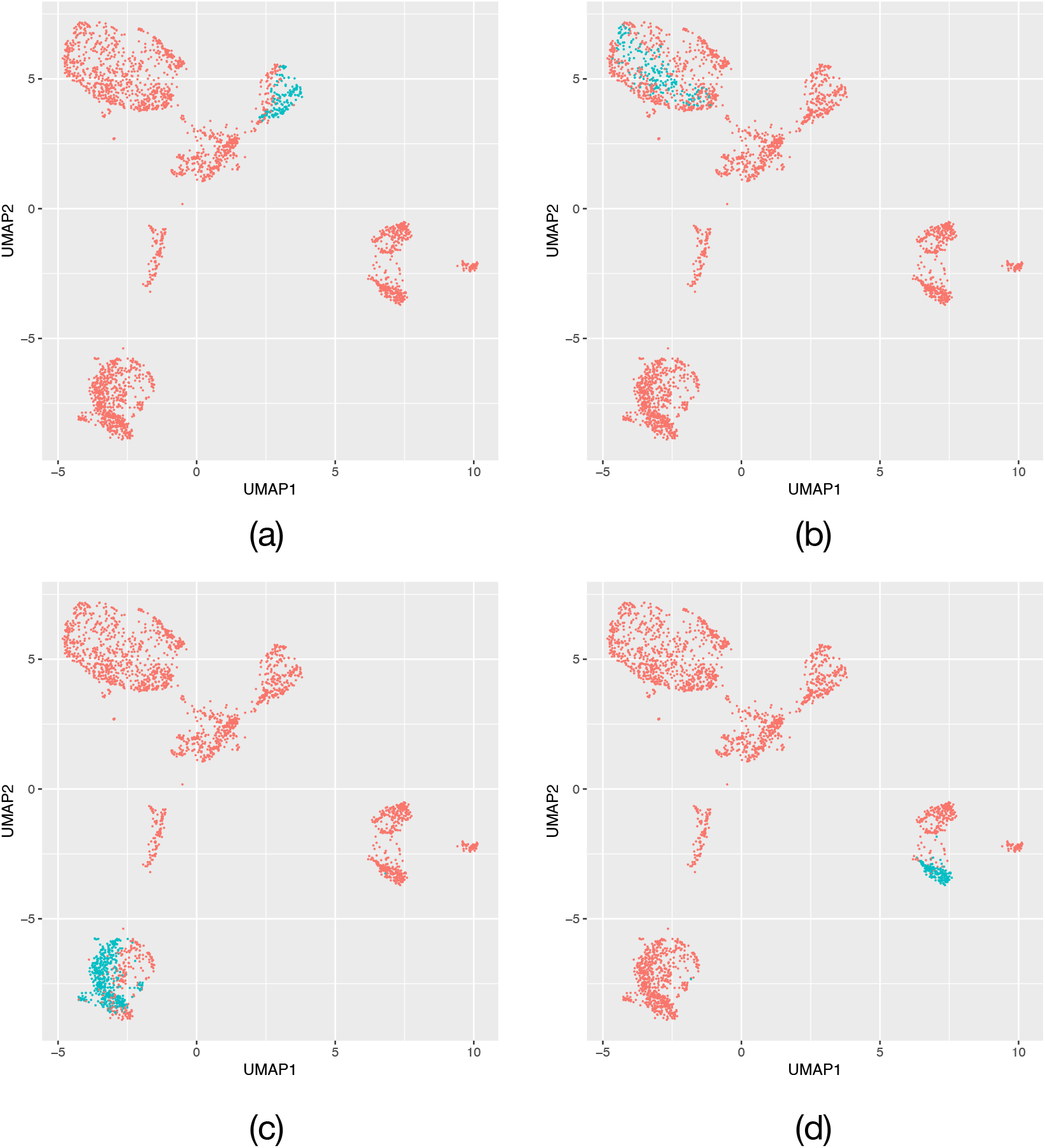
UMAP embeddings of cells from the mouse cortex and hippocampus data Zeisel et al. (2015) with clusters (a) 1, (b) 7, (c) 18 and (d) 20 identified by our method highlighted in blue.

**Figure 6:**
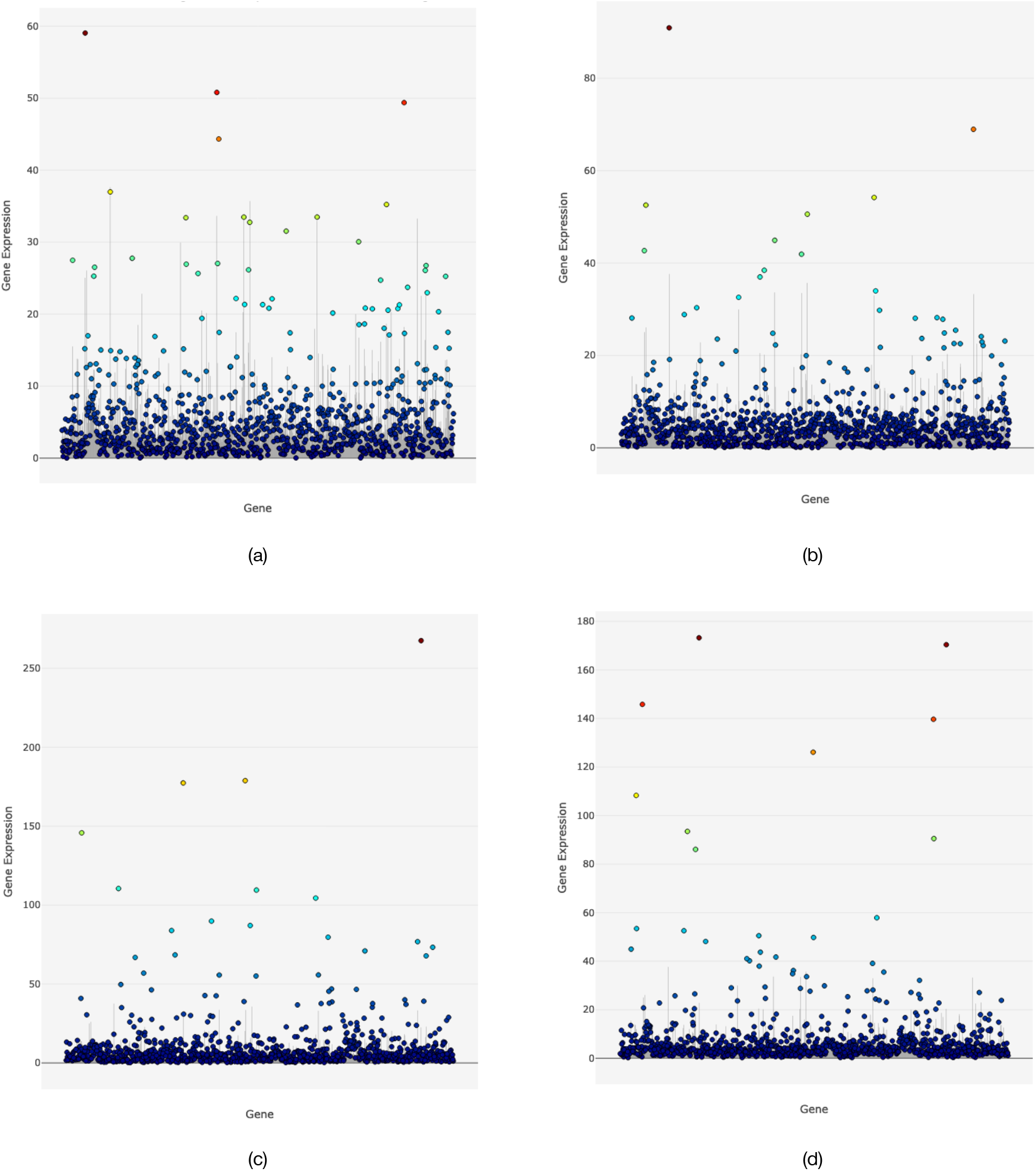
Cluster specific means in the zero-inflated negative binomial model for each gene inferred from the mouse cortex and hippocampus data Zeisel et al. (2015) for clusters (a) 1, (b) 7, (c) 18 and (d) 20. Cluster means are shown as coloured circles and the average expression across all clusters shown as grey bars.

In cluster 1, upregulated genes include *Gad1, Gad2* and *Slc6a1*, known markers of interneurons. Looking at the set of upregulated genes in cluster 1, it was found to be enriched for genes in the transmission across chemical synapses pathway, and the GABA synthesis, release, reuptake and degradation pathway, amongst others. The enrichment of genes in the GABA synthesis pathway may indicate that this specific cluster represents GABAergic interneurons, that synthesise the gamma-aminobutyric acid (GABA) neuro-transmitter.

Cluster 7 corresponds to pyramidal CA1 neurons, and in the gene set enrichment analysis we found that upregulated genes for this cluster were enriched for genes in the AMPA receptors pathway. Pyramidal neurons in the hippocampus respond to excitatory glutamate via AMPA receptors, and the AMPA receptors *Gria1* and *Gria2* are both upregulated in this cluster. In cluster 18 on the other hand oligodendrocyte specific gene *Mog* and the gene *Mbp* that encodes a protein that is a major component of the myelin sheath, are discovered to be upregulated. Oligodendrocytes play a central role in the formation and maintenance of the myelin sheath that covers neurons (Simons & Nave 2016). The set of upregaulted genes in cluster 18 was also enriched for genes in the axonal guidance and nervous system development pathways, and it is known that oligodendrocytes inhibit the growth of axons as they develop (Fawcett et al. 1989).

Several marker genes of astrocytes are found to be upregulated in cluster 20, including *Slc1a2, Slc1a3, Apoe, Glul* and *Aqp4*. Considering the set of upregulated genes, cluster 20 is enriched for genes in the glutamatergic synapse pathway, and astrocytes express glutamate (an excitatory neurotransmitter) uptake transporters (Sofroniew & Vinters 2010). One of the overexpressed genes, *Glul*, encodes an astrocyte specific protein that plays a central role in the uptake of glutamate at synapses.

## 4 Discussion

In this paper we have introduced a novel Bayesian nonparametric zero-inflated negative binomial model for the clustering of scRNA-seq data, comprised of a mixture model at the count level for individual genes to allow for variation in the degree of zero-inflation, as well as a second mixture at the cell level, and demonstrated that it can perform better than SC3 when applied to many of the data sets in the benchmarking data used, although Seurat still outperforms our model on the majority of these. Results on benchmarking data also demonstrate that the zero-inflated negative binomial model performs better than the negative binomial and multinomial probabilistic models we considered.

Both the Seurat and SC3 software packages allow for the identification of differentially expressed or marker genes within clusters, by applying existing differential expression methods to the cells within a cluster, in Seurat and SC3, or by training a classifier on expression within clusters with SC3. These approaches would be equally applicable to results from our methods, with the added benefit of the inclusion of uncertainty in cluster membership.

A benefit of our approach is that we can derive a statistical model of gene expression within each sub-population of cells identified. This allows us to easily identify genes that are over or under expressed in a sub-population of cells without post-hoc examination of the clustering. However the key benefit of our approach is that by applying a Bayesian model we are also able to quantify the uncertainty in cluster allocation for each cell. Furthermore we demonstrate that the zero-inflated negative binomial model we propose outperforms the multinomial and negative binomial Dirichlet process mixture models of scRNA-seq counts in identifying clusters and finding the correct number of clusters on the benchmarking data used. In the supplementary material, through the use of statistical model selection we show that for some genes a zero-inflated model is preferable to a negative binomial.

Applying the method to real world scRNA-seq data we are able to identify clusters of cells corresponding to cell types identified in previous work in the literature, and find cluster specific overexpressed genes that correspond to known cell type markers. We also identify enriched pathways with known functional relevance to the cell types considered in the sets of upregulated genes.

We see this methodology as a building block for scRNA-seq data analysis pipelines that are able to report and propagate uncertainty (Lähnemann et al. 2020), as cell type clustering is often one of the first steps applied before performing downstream analyses such as differential expression (Kharchenko et al. 2014, Ntranos et al. 2019), or network inference (Pratapa et al. 2020, Stumpf 2021). To allow Gaussian Graphical Models (GGMs), which have been used extensively in the modelling of gene regulatory networks (Scutari & Strimmer 2011), to be applied to scRNA-seq data, copula approaches that transform the marginal cumulative distributions of the data to match those of a Gaussian distribution could be applied. This could make use of the inferred within cluster distributions for the distribution of counts for each gene to use in the copula transformation.

A benefit of our Bayesian nonparametric approach is that the prior adapts naturally to the complexity of the data and can represent an unbounded number of clusters as the size or heterogeneity of the data grows. Running times on a single CPU were on the order of 5 hours on a data set of around 3000 cells, and in future work we will consider approaches to accelerating the inference using approximate Bayesian methods such as stochastic variational inference (Hoffman et al. 2013), that will allow this approach to scale to larger data sets of potentially millions of cells.

## Acknowledgements

We would like to thank the anonymous reviewers for their detailed and helpful comments that enabled us to improve the readability and presentation of the manuscript.

## Availability of data and software

The single cell benchmarking data from Tian et al. (2019) are available from Gene Expression Omnibus https://www.ncbi.nlm.nih.gov/geo/ with accession GSE118767, and from https://github.com/LuyiTian/sc_mixology. The mouse cortex and hippocampus data from Zeisel et al. (2015) are available from GEO with accession GSE60361 and from https://linnarssonlab.org/cortex/. The code used to produce the results shown in the manuscript is available as a Snakemake pipeline(Köster & Rahmann 2012) from https://github.com/tt104/scmixture. The Snakemake pipeline is also archived along with a Singularity container to run the code on Zenodo, DOI 10.5281/zen-odo.6639073.

## Supplementary information

### Model selection of within-cluster distributions

We consider two possible models for the observed scRNA-seq counts *y*_*ij*_ of a single gene *j* within a single cell type. These models are a negative binomial distribution

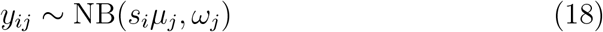

where *s*_*i*_ is a scaling factor to account for sequencing depth, *μ*_*j*_ is the mean and *ω*_*j*_ the dispersion. We also consider a zero-inflated negative binomial distribution

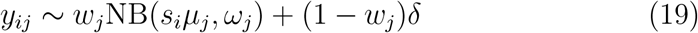

which has a further parameter *w*, determining the rate of dropout of the gene in a given cell type. Here *δ* represents the Dirac delta distribution.

To investigate the appropriate model for single cell RNA-seq counts within a given cell type, we applied the Bayesian Information Criterion (BIC) given in equation 20 to maximum likelihood parameter estimates 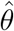 for individual genes from a subset of 1000 selected as having the largest coefficient of variation. We applied analysis to this subset of genes as clustering is typically applied to a subset of the genes in the data having the largest variance (Duan et al. 2019), and so these are the genes for which it is important to identify whether zero-inflation is present and should be included in our model.

We compared the models of a negative binomial (equation 18) and zero-inflated negative binomial (equation 19) distribution for the likelihood function *f*, using the BIC

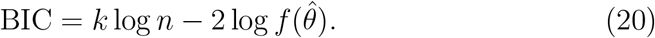

Here *k* represents the number of parameters of the model, and penalises unnecessarily complex models, while *n* is the number of data points observed. Using the model selection approach described above, we considered the best models for count data from scRNA-seq benchmarking experimental data(Tian et al. 2019) as well as an scRNA-Seq data set from the mouse cortex and hippocampus taken from the literature(Zeisel et al. 2015).

Results are presented in figure S1, showing the percentage of genes in each cell type within an experiment that were found to be better explained by the zero-inflated model. It is apparent that there are zero-inflated genes across each data set and cell type, and although the percentages are low, they are consistent across experimental platforms. This suggests that using a model that is able to include zero-inflation at different degrees across each gene is a more realistic model of real world scRNA-seq data than using a negative binomial model alone. Further when considering the maximum likelihood estimates for dropout rates in each cell type, we see that there is variation in dropout rate across the different cell types in figure S2. This motivates the choice of a cluster specific dropout rate for each gene.

**Figure S1:**
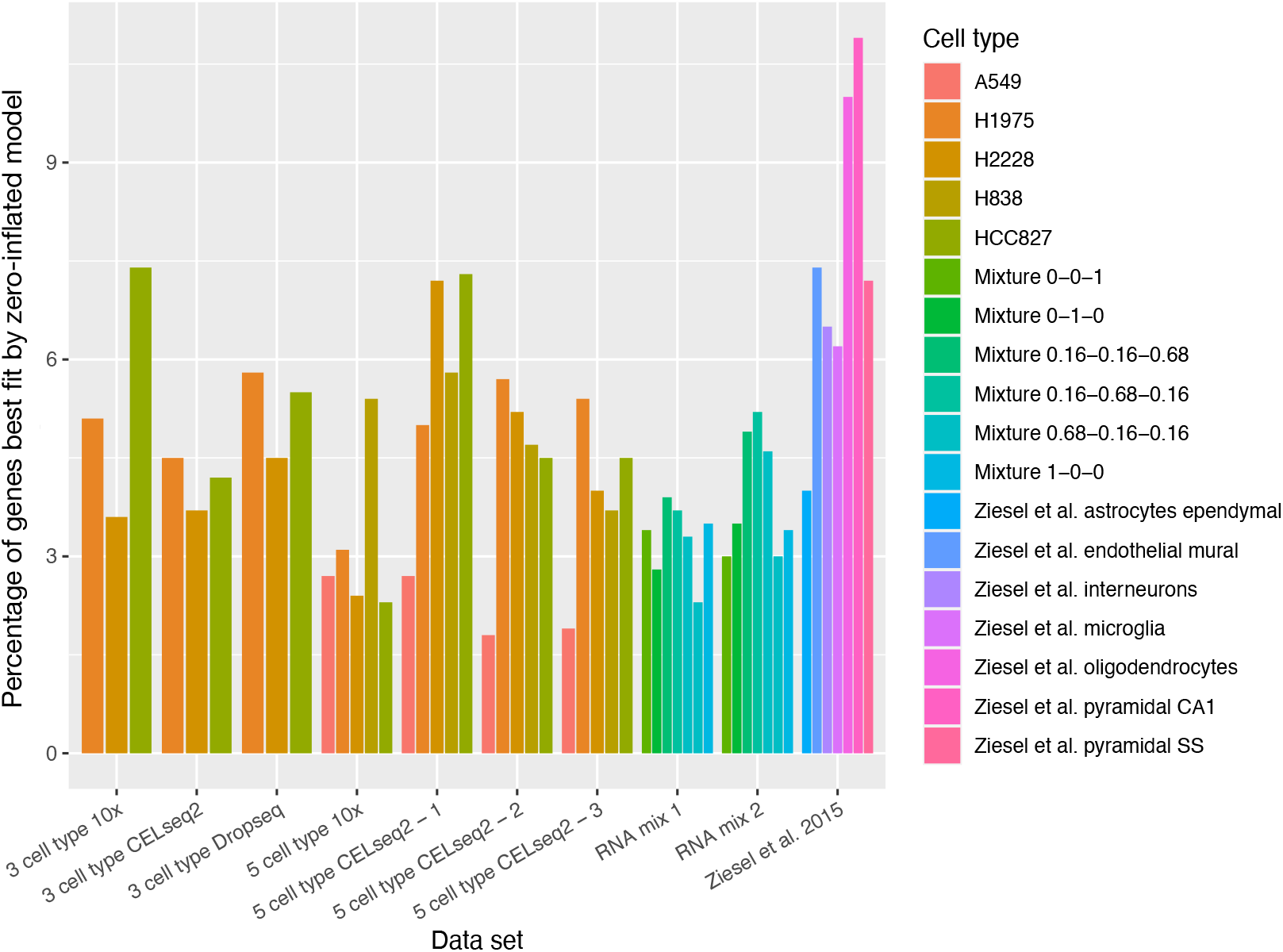
Percentages of genes in each cell type, across data sets from Tian et al. (2019) (data sets 1-9) and Zeisel et al. (2015), where the zero-inflated negative binomial model is preferred to the negative binomial model under the BIC.

**Figure S2:**
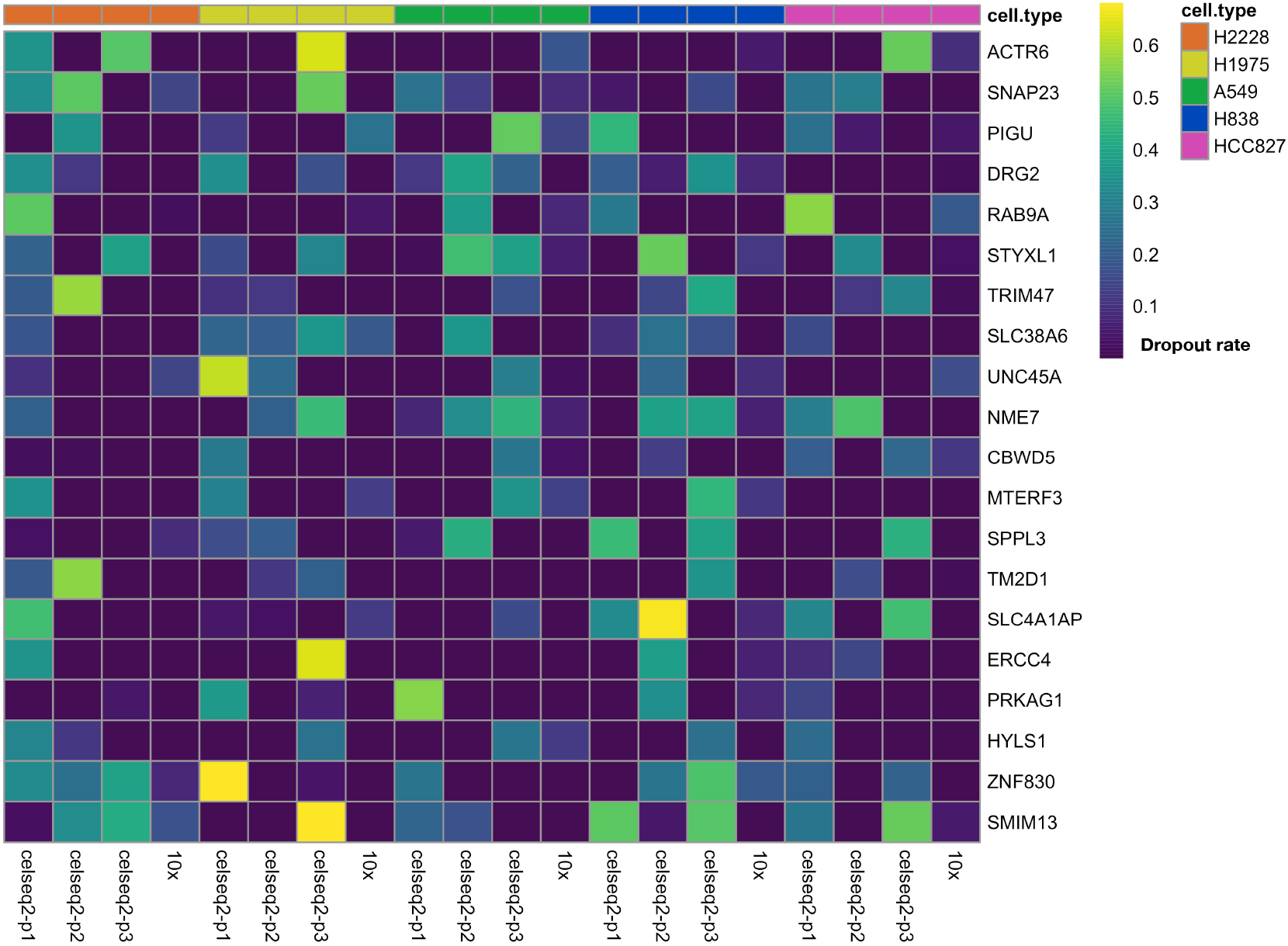
Heatmap showing variation in the inferred dropout rate parameter for a zero-inflated negative binomial model for a selection of genes across several experimental platforms and cell types from Tian et al. (2019). It can be seen that for a single gene (rows) the inferred dropout rate of the zero-inflated negative binomial model varies between cell types (grouped columns annotated at the top of the figure) and experimental platforms (columns).

